# Explainable AI reveals the quantitative hierarchical architecture of global bird extinction risk

**DOI:** 10.64898/2026.05.18.726070

**Authors:** Pablo Medrano-Vizcaíno, Atriya Sen, Anthony Marchiafava

## Abstract

Identifying what makes species vulnerable to extinction requires accounting for complex biological and environmental interactions. Due to their high predictive accuracy, machine learning methods have been widely used for these assessments; however, relying on black-box models offers limited interpretability. Here, using a comprehensive dataset of anthropogenic, ecological, morphological, demographic, and biogeographical variables from 9,053 species (81% of birds worldwide), we applied Inductive Logic Programming (ILP), an explainable artificial intelligence framework, to generate explicit and quantitative IF-THEN rules with confidence scores for bird extinction risk. Our approach revealed that extinction vulnerability follows a hierarchical structure, shaped by interactions among range size, morphological traits, and human pressures. The framework recovered well-established knowledge, while also revealing previously undescribed extinction patterns. For example, consistent with prior evidence, species with geographic ranges below ∼13,500 km² were identified as higher risk (88% confidence). Nevertheless, this threshold shifted to ∼3,270 km² when human impacts were removed, revealing quantitatively how anthropogenic activities expand the pool of vulnerable species beyond those at risk due to biological and biogeographical traits alone. Beyond established patterns, species with tail length >304 mm were identified as higher risk (82% confidence), a pattern not previously documented. ILP models achieved 91% overall accuracy, slightly lower than Random Forest (93%), but notably better than Neural Networks (83%). These results show that ILP can offer high accuracy results with full interpretability, also providing quantitative transition thresholds that clarify the structural architecture of extinction risk, and translate complex ecological interactions into actionable tools for conservation.

## 1. Introduction

During the last century extinction rates have increased dramatically, becoming 10-100 times higher than the natural average background rates (Ceballos et al. 2015). Although natural causes can be involved as driving factors, human activities associated with land-use change, resource exploitation, pollution, climate change and invasive species, have been identified as the main causes (Keck et al. 2025). Interestingly, the magnitude of effects differs among species depending on their ecology, morphology, demography and geographic distribution (Purvis et al. 2000; Chichorro et al. 2022). These aspects can define the response of species to human disturbances and their vulnerability to extinction (Purvis et al. 2000).

Several studies have found that large species with a more specialized diet and occupying less habitats are more vulnerable to extinction (Purvis et al. 2000; González-Suárez et al. 2013; Martínez et al. 2023). A similar trend has been found for species with a smaller range size, and a narrower altitudinal breadth (Purvis et al. 2000; Keane et al. 2005), highlighting the importance of biogeographical variables. This is also the case of long-lived species with a low population density, and an increased generation length (Bestion et al. 2015; Bird et al. 2020), demonstrating the role of demographics.

Collectively, these findings indicate that extinction risk emerges from combinations of ecological, biogeographical, and anthropogenic factors that interact in complex ways. However, most research has examined these drivers in isolation (i.e., ecological, morphological, and human pressure data analyzed separately) overlooking their synergistic effects (Tobias et al. 2022).

Understanding how multiple processes combine to shape extinction risk is essential for gaining a holistic perspective on the interplay between species traits, environments, and human pressures. Although a few papers have considered this holistic perspective, and have revealed relevant findings for conservation purposes (González-Suárez et al., 2013; Stewart et al., 2025), merely finding statistical correlations between various factors is not sufficient for effective conservation, which requires an explicit understanding of how extinction patterns are structured.

Birds represent an ideal model taxon for investigating global extinction patterns at a planetary scale. They comprise the most comprehensively assessed vertebrate class in the IUCN Red List (>11,000 extant species with detailed threat classifications), and have extensive trait databases available (Tobias et al. 2022; Şekercioğlu et al. 2025), enabling unprecedented integration of ecological, life-history, biogeographical, and anthropogenic drivers in a single analytical framework. Nevertheless, this data may require high complexity analyses, an area where sophisticated tools such as machine learning, and deep learning techniques excel due to their capacity to deal with big and multidimensional data, and to outperform traditional statistical models in predictive accuracy (Cutler et al., 2007). Indeed, machine learning techniques (e.g., Random Forests and Neural Networks) have been increasingly adopted across a wide range of conservation applications, such as species distribution modelling, population trend prediction, and extinction risk classification (Branco et al., 2023). Despite advantages, these tools are considered as black-box models, due to their lack of interpretability.

In contrast to black-box models, Inductive Logic Programming (ILP), a subfield of symbolic artificial intelligence that combines machine learning and logic programming, provides an effective framework for generating transparent and explainable rules (Cropper et al. 2022). ILP systems learn by constructing logical rules from data, producing outputs in the form of “IF-THEN” statements that explicitly describe the conditions leading to specific outcomes (Mcginness & Baumgartner 2024).

Despite its potential, ILP has been rarely applied to large-scale ecological problems (but see Tamaddoni-Nezhad et al. 2012; Tamaddoni-Nezhad et al. 2014). This gap stems from the fact that traditional ILP systems have faced significant challenges handling numerical data (requiring discretization that loses information), restriction to binary outputs rather than multiclass predictions, and limited scalability for big datasets (Ielo et al. 2023; Cropper et al. 2025). These constraints have prevented their adoption in biodiversity science where datasets are inherently large, complex, and computationally demanding.

However, recent ILP systems have been capable of overcoming these limitations (Mcginness & Baumgartner 2024), for example incorporating numerical optimization techniques, supporting multiclass classification through predicate hierarchies, and leveraging parallel computing architectures to handle large datasets. For instance, rather than providing a probabilistic extinction risk score without explanation, ILP can generate rules such as: “IF a species has a range size ≤5,000 km², a body mass of ≥100 g, AND a tarsus length ≤25 mm THEN extinction risk is high (Confidence: 80%)”. Note that these rules explicitly contain numerical thresholds, combine features and provide a confidence score, which evidently is a step forward large-scale analysis that have relied on traditional statistical approaches or black-box models such as Neural Networks. These advances, combined with pruning strategies that reduce search space complexity, make it feasible for the first time to apply ILP to comprehensive global biodiversity assessments while maintaining the interpretability that has always been ILP’s core strength. This is fundamental for conservation science, where understanding why a species is threatened is as important as predicting that it is threatened.

Here we leverage these recent advances in ILP using a comprehensive global dataset of ecological, morphological, biogeographical, and anthropogenic variables for >9,000 bird species, and derive the first fully explainable, quantitative rule-based assessment (with confidence scores) of avian extinction risk. Finally, we compare ILP performance against typical black-box models (Random Forest, Neural Networks) and demonstrate that high explainability and high predictive power can be achieved for large-scale analyses.

## 2. Methods

### 2.1 Data management

#### 2.1.1 Variables selection

We compiled species-level data from multiple global datasets covering different types of variables: biogeographical (Tobias et al. 2022; Şekercioğlu et al. 2025), demographic (Bird et al. 2020), ecological (Tobias & Pigot 2019; Tobias et al. 2022; Şekercioğlu et al. 2025), morphological (Tobias et al. 2022; Şekercioğlu et al. 2025), and human threats. Human-threat variables were obtained using the rrdelist (Gearty & Chamberlain 2025) R package, which provides species-level data directly from the IUCN Red List API. We also included the taxonomic order and family of each species to account for evolutionary relatedness and for similarities across taxa that may not be captured by the measured traits.

To reduce redundancy among predictors, we computed a correlation matrix and retained only variables with r<0.75 (Figure S1), except in cases where correlated variables represented biologically distinct dimensions. For example, tarsus length and wing length (r=0.76) were both retained because they relate to terrestrial locomotion and flight performance, respectively. Similarly, elevational range and maximum altitude (r=0.79) were retained as elevational range reflects the breadth of altitudes a species occupies, whereas maximum altitude indicates the maximum altitude at which a species is known to occur. Also, Generation length and wing length (r=0.78) were retained as they represent different demographic and morphology aspects.

#### 2.1.2 Extinction risk

We used the IUCN Red List conservation status as a proxy to study extinction risk. This represents the most accepted measure of extinction risk globally, and its use is widespread in both scientific and policy contexts (Mace et al., 2008). Each species was assigned its current IUCN Red List category (LC–Least Concern, NT–Near Threatened, VU–Vulnerable, EN–Endangered, CR–Critically Endangered) (https://www.iucnredlist.org). Later, all species were grouped into two classes: Lower risk (LC) and Higher risk (NT, VU, EN, CR). We did this classification to reduce the imbalance among categories, an approach previously used in other studies (Ali et al. 2023). DD-Data deficient species were not included in any class. Finally, our dataset included 9,053 species (1,722 Higher risk species and 7,331 Lower risk species) (i.e., 81.2% of all known bird species in the planet).

#### 2.1.3 Taxonomic backbone

Datasets can have different taxonomic nomenclatures, therefore, scientific names from all datasets were standardized to the HBW/BirdLife Taxonomic Checklist (version 10) (https://datazone.birdlife.org/about-our-science/taxonomy), which is the standard taxonomy for global bird assessments such as IUCN Red List evaluations, Important Bird and Biodiversity Areas (IBAs), and Key Biodiversity Areas (KBAs).

Each species name was queried against the GBIF API to retrieve the currently accepted name along with synonyms. Trait databases were then integrated through a progressive matching procedure, attempting matches sequentially using the backbone name, the GBIF accepted name, and synonyms, maximizing taxonomic coverage and trait-data completeness across sources.

### 2.2 Data Analysis

We compared three classification approaches to predict extinction risk: an Explainable AI method based on Inductive Logic Programming (CON-FOLD), Random Forest, and a Neural Network. We did this to evaluate the consistency of results across methods with different levels of interpretability and complexity and to assess whether the predictive performance of ILP was comparable to well-established machine learning methods. All models were evaluated using 10-fold cross-validation, where data partitions were created using the createFolds function from the caret package in R (Kuhn, 2008). The same folds were applied consistently across all three methods to ensure fair and direct comparison. For each fold, precision, recall, F1-score, and overall accuracy were calculated. Performance estimates are reported as the mean and standard deviation across folds.

#### 2.2.1 Explainable AI through ILP

We used CON-FOLD (Mcginness & Baumgartner, 2024), an explainable machine learning classification algorithm that generates hierarchical rule sets in a sequential decision structure with associated confidence scores. This system is highly scalable, can handle numerical and noisy data, and allows the integration of expert knowledge (Mcginness & Baumgartner, 2024). We applied a pruning threshold of 20% using the confidence_fit(…) method, which means that exceptions to the rules are added only if they improve rules’ confidence by that specified threshold, thereby avoiding unnecessary complexity. In addition, we kept only higher confidence rules (≥75%).

We generated two types of models: 1) Data-driven models, where CON-FOLD induces the rules directly from the analysis of the dataset, and 2) Expert models, which include expert rules (also known as background knowledge, a key advantage of ILP) with a confidence value representing the certainty with which we expect the rule to hold.

In addition, because human threats are known to be strong predictors of extinction risk (Chichorro et al. 2019; Dueñas et al. 2021; Hinsley et al. 2023; Cabernard et al. 2024), each type of model (data-driven and expert) had two types of submodels: one including human threat variables, and another without them. This was done to isolate the explanatory contribution of species traits and other extrinsic factors, while minimizing potential circularity arising from the use of IUCN threats to predict extinction risk based on IUCN categories.

##### Expert rules

These rules were based on well-documented extinction-risk patterns found in scientific literature: species with small geographic ranges and whose populations are heavily affected by agriculture, invasive species, and hunting tend to be more vulnerable to extinction (Chichorro et al. 2019; Dueñas et al. 2021; Hinsley et al. 2023; Cabernard et al. 2024). The same has been reported for large species with long generation length and narrow altitudinal range (Bird et al. 2020; Chichorro et al. 2022). In addition, we also relied on feature importance rankings from Random Forest analyses for identifying the most important variables for rules construction (see the Random Forest methods section for details on this calculation).

Therefore, we classified our empirical data in quartiles, where the first quartile was related to “small” or “low” whereas values in the third quartile were classified as “large” or “long”. Then, thresholds used in the following expert rules correspond to the empirical quartile boundaries derived from the dataset.

Expert models with human threats included this background knowledge (confidence score is shown in brackets at the end of each rule):

- small range size (≤75,321 km²) and affected by agriculture, hunting, and invasive species → higher risk (99%).
- affected by agriculture, hunting, and invasive species → higher risk (95%).
- small range size (≤75,321 km²) and affected by hunting and invasive species →higher risk (95%).
- small range size (≤75,321 km²) and affected by agriculture and hunting → higher risk (95%).
- small range size (≤75,321 km²) and affected by agriculture and invasive species→ higher extinction risk (95%).

While expert models without human threats included this background knowledge:

- Large species (>130 g), small range size (≤75,321 km²), long generation length (>4.068 years) and narrow altitudinal range (≤800 m) → higher risk (99%).
- Large species (>130 g), small range size (≤75,321 km²), and narrow altitudinal range (≤800 m) → higher risk (95%).
- small range size (≤75,321 km²), long generation length (>4.068 years) and narrow elevational range (≤800 m) → higher risk (95%).
- Large species (>130 g), small range size (≤75,321 km²), and long generation length (>4.068 years) → higher risk (95%).
- Large species (>130 g), long generation length (>4.068 years) and narrow elevational range (≤800 m) → higher risk (95%).

#### 2.2.2 Random Forest

This machine learning method, which generates an ensemble of multiple classification trees from a bootstrap sample of the original data, is known for highly accurate predictions, surpassing other methods such as logistic regression and linear discriminant analysis (Cutler et al. 2007). Random forests were built with 2,000 trees using the “ranger” function from the R package ranger (Wright & Ziegler 2017). Each model incorporated all trait variables, taxonomic order, family and spatial predictors such as minimum latitude and longitude of the species distribution to broadly represent spatial variation. Variable importance was estimated using the function importance () from the R package ranger (Wright & Ziegler 2017), based on the permutation importance metric. Also, we generated partial dependence plots using the function partial () from the R package pdp (Greenwell 2017) to visualize the relationships between predictor variables and the predicted extinction risk.

#### 2.2.3 Neural networks

Neural Networks are powerful general function approximators that learn weights to map inputs to outputs (Hastie et al., 2009). We applied the IUCNN model (Zizka et al., 2022) as the most relevant point of comparison. IUCNN supports multiple task types: a regression model to predict specific values, a convolutional neural network for image-based classification, and a fully connected feed-forward network for classification on numerical inputs. The latter was selected due to its performance at classifying IUCN status and its use of similar input data to that used here. Because the input features differ between the original implementation and the present study, parts of the network architecture were modified accordingly.

## 3. Results

Our compiled dataset included complete traits for 9,053 species out of the 11,149 total species (81.2% of birds globally). This included 31 variables: 6 ecological, 7 morphological, 9 biogeographical, 3 demographic, 2 taxonomic, and 4 related to human pressures.

All three approaches (CON-FOLD, Random Forest, and Neural Networks) achieved overall accuracy above 80%. The best-performing model was Random Forest including human threats, achieving 93%, followed by CON-FOLD expert (90.56%) and CON-FOLD data-driven (89.24%) (Table 4). However, performance metrics were consistently higher when predicting the lower-risk class, likely reflecting class imbalance (7,331 Lower risk species vs 1,722 Higher risk species). The inclusion of human threat variables improved all models’ performance, particularly for the higher-risk class.

**Table 1.** Variables used to predict extinction risk and their definitions.

| Type | Variable | Definition | Source |
| --- | --- | --- | --- |
| Biogeographic | Range size | The total area of the mapped range of the species. | Tobias et al., 2022 |
|  | Minimum elevation | Normal lowest elevational limit of species | Şekercioğlu et al., 2025 |
|  | Maximum elevation | Normal highest elevational limit of species | Şekercioğlu et al., 2025 |
|  | Elevational range | The difference between a species' normal lowest and highest elevational extents. | Şekercioğlu et al., 2025 |
|  | Minimum latitude | The minimum latitudinal extent of the species range (restricted to breeding and resident range) | Tobias et al., 2022 |
|  | Maximum latitude | The maximum latitudinal extent of the species range (restricted to breeding and resident range) | Tobias et al., 2022 |
|  | Latitudinal range | Latitudinal range of species based on tropics and polar circles: 1: Tropical, 2: Tropical-Temperate, 3: Temperate, 4: Temperate-Polar, 5: Tropical-Polar | Şekercioğlu et al., 2025 |
|  | Realm | Biogeographical realms: large-scale regions defined by shared evolutionary history, distributional patterns and distinctive species assemblages. A: Australian, C: Cosmopolitan, E: Eastern Hemisphere, F: Afrotropical, I: Indomalayan, L: Neotropical, M: Madagascar & islands, N: Nearctic, O: Oceania, P: Palearctic, S: South Polar, W: Wallacea, Z: New Zealand & islands | Şekercioğlu et al., 2025 |
|  | Island restricted breeding | Yes or No | Tobias & Pigot et al., 2019 |
| Demographic | Adult survival | Probability of annual adult survival | Bird et al., 2020 |
|  | Generation length | Average age of parents of the current cohort (i.e. newborn individuals in the population), reflecting the turnover rate of breeding individuals in a population. | Bird et al., 2020 |
|  | Clutch size | Number of offspring per reproductive event | Tobias & Pigot et al., 2019; Şekercioğlu et al., 2025 |
| Ecological | Habitat category | Artificial = Artificial, agricultural, farms, plantations, suburban, quarry, mine, excavation, dams and man-made lakes.<br>Bamboo = Bamboo lands.<br>Coastal = intertidal zones within immediate vicinity of beaches, estuaries, brackish to salty marshes;<br>Desert = drylands and other open arid habitats, often sandy with very sparse vegetation;<br>Forest = tall tree-dominated vegetation;<br>Grassland = open dry to moist grass-dominated landscapes;<br>Plains = Plains, dry & open areas, semi-desert, steppe, tundra, arid and semi-arid grassland.<br>Riparian = Riparian, riverine, gallery forest, stream, running water, wooded ravines.<br>Rocky = rocky substrate typically with no or very little vegetation;<br>Savanna = Savanna, arid plains, wooded grassland, pampas;<br>Sea = Open sea, pelagic;<br>Shrub = low stature bushy habitats;<br>Wetland = lakes, marshes, swamps and reedbeds;<br>Woodland = medium stature tree-dominated habitats. | Tobias et al., 2022; Şekercioğlu et al., 2025 |
|  | Habitat breadth | Habitat breadth, the number of major habitats used | Şekercioğlu et al., 2025 |
|  | Diet | Frugivore = at least 60% of diet from fruit;<br>Granivore = at least 60% diet from seeds or nuts;<br>Nectarivore = at least 60% diet from nectar;<br>Herbivore = at least 60% diet from other non-aquatic plant materials;<br>Herbivore aquatic = at least 60% of diet from aquatic plants;<br>Invertivore = at least 60% of diet from terrestrial invertebrates;<br>Vertivore = at least 60% of diet from terrestrial vertebrates; | Tobias & Pigot et al., 2019; Tobias et al., 2022; Şekercioğlu et al., 2025 |
|  |  | Aquatic animals = primary diet from aquatic vertebrates and invertebrates;<br>Beeswax = Primarily feeds on bees wax;<br>Scavenger = at least 60% of diet from carrion, offal or refuse;<br>Omnivore = using multiple niches, within or across trophic levels, in relatively equal proportions |  |
|  | Diet breadth | Diet breadth, the number of major food types consumed | Şekercioğlu et al., 2025 |
|  | Migration | Sedentary, partially migratory (minority of population migrates long distance or most individuals migrate short distances) and migratory (majority of population undertakes long-distance migration) | Tobias & Pigot et al., 2019, Şekercioğlu et al., 2025 |
|  | Primary lifestyle | Aerial = much of the time in flight;<br>Terrestrial = majority of time on the ground;<br>Insessorial = much of the time perching above the ground;<br>Aquatic = much of the time on water;<br>Generalist = spends time in different lifestyle classes | Tobias et al., 2022 |
| Human threats | Agriculture | The population of x species is affected by this pressure (1) or not (0). | IUCN red list |
|  | Climate change | The population of x species is affected by this pressure (1) or not (0). | IUCN red list |
|  | Hunting | The population of x species is affected by this pressure (1) or not (0). | IUCN red list |
|  | Invasive species | The population of x species is affected by this pressure (1) or not (0). | IUCN red list |
| Morphological | Body mass | Average body mass (adult males and females). | Tobias et al., 2022 |
|  | Tarsus length | Length of the tarsus from the posterior notch between tibia and tarsus, to the end of the last scale of acrotarsium (bend of the foot) | Tobias et al., 2022 |
|  | Wing length | Length from the carpal joint (bend of the wing) to the tip of the longest primary on the unflattened wing | Tobias et al., 2022 |
|  | Beak depth | Depth of the beak at the anterior edge of the nostrils | Tobias et al., 2022 |
|  | Tail length | Distance between the tip of the longest rectrix and the point at which the two central rectrices protrude from the skin, typically measured using a ruler inserted between the two central rectrices | Tobias et al., 2022 |
|  | Beak length_culmen | Length from the tip of the beak to the base of the skull | Tobias et al., 2022 |
|  | Hand wing index | 100*DK/Lw, where DK is Kipp's distance and Lw is wing length (i.e., Kipp's distance corrected for wing size). | Tobias et al., 2022 |
| Taxonomic | Family | Taxonomic family | Birdlife |
|  | Order | Taxonomic order | Birdlife |

**Table 2.** Importance of predictors from the Random Forest model including human threats.

| Variable | Importance |
| --- | --- |
| Agriculture | 0.062 |
| Hunting | 0.054 |
| Range size | 0.029 |
| Invasive species | 0.028 |
| Maximum latitude | 0.007 |
| Climate change | 0.006 |
| Generation length | 0.006 |
| Minimum latitude | 0.006 |
| Elevational range | 0.005 |
| Wing length | 0.005 |
| Body mass | 0.005 |
| Maximum elevation | 0.004 |
| Beak depth | 0.004 |
| Island restricted breeding | 0.004 |
| Adult survival annual | 0.003 |
| Tarsus length | 0.003 |
| Beak length culmen | 0.003 |
| Realm | 0.003 |
| Tail length | 0.002 |
| Hand wing index | 0.002 |
| Minimum elevation | 0.002 |
| Clutch size | 0.002 |
| Habitat breadth | 0.001 |
| Family | 0.001 |
| Diet | 0.001 |
| Latitudinal range | 0.001 |
| Order | 0.001 |
| Habitat | 0.001 |
| Primary lifestyle | 0.001 |
| Migration | 0.001 |
| Diet breadth | 0.000 |

**Table 3.**
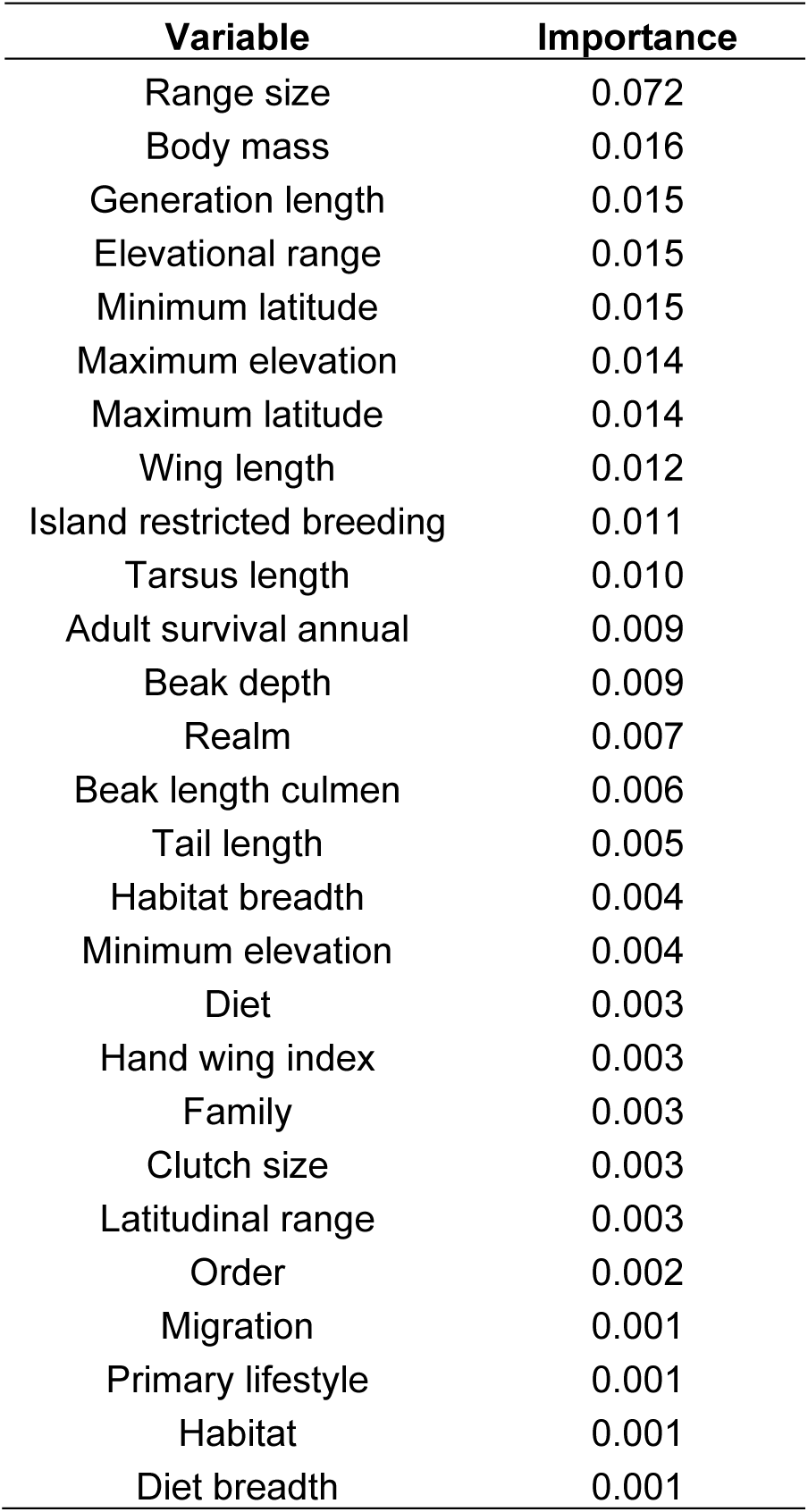
Importance of predictors from the Random Forest model not including human threats.

**Table 4.**
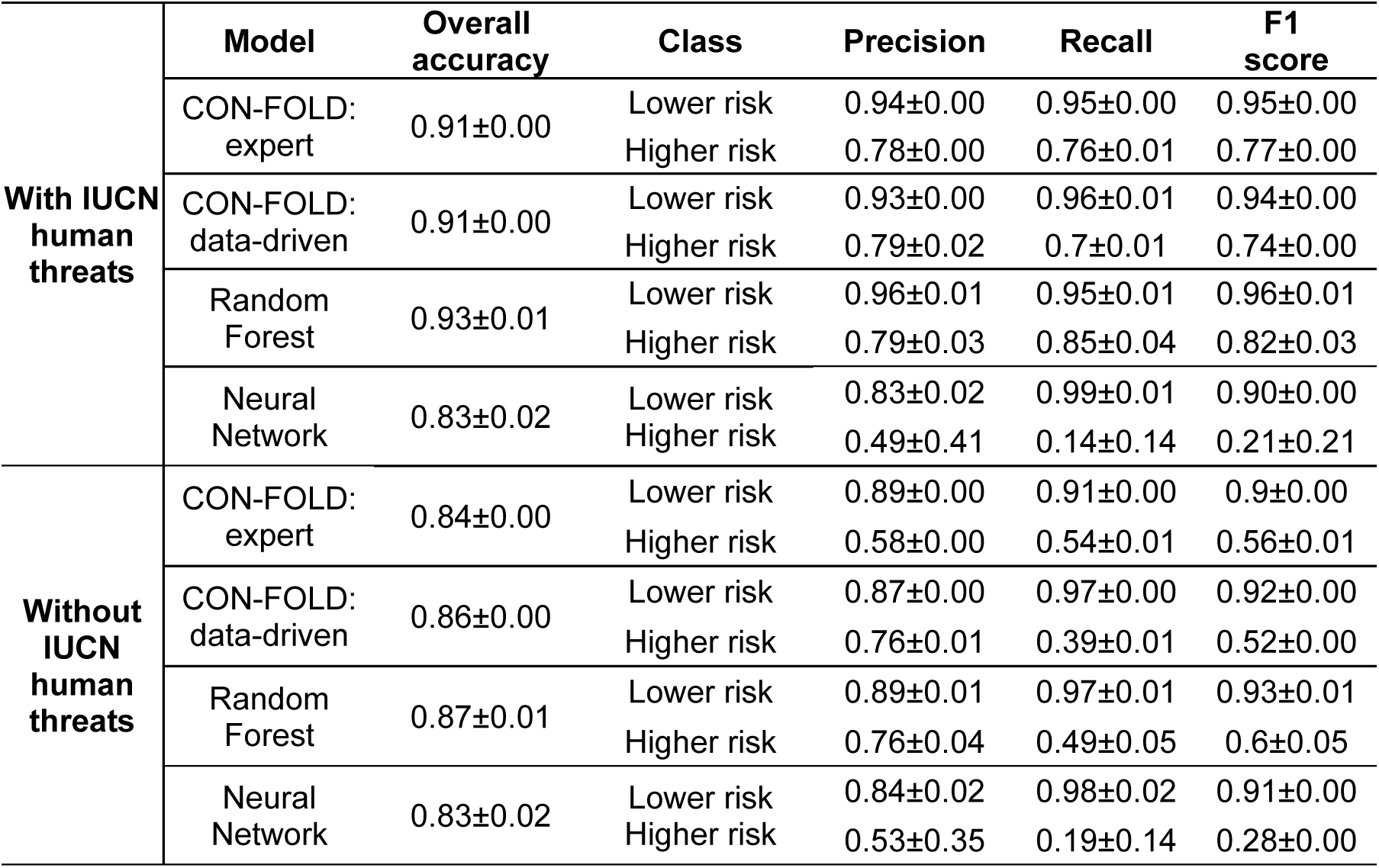
Metrics comparison between CON-FOLD, Random Forest and Neural Network models with and without IUCN human threats (mean ± SD across 10 stratified folds).

Among human pressures, agriculture emerged as the most important predictor (positively associated with extinction risk). Nevertheless, biological and biogeographical traits also revealed strong extinction patterns, with range size and elevational range consistently ranking among the top predictors across models.

### 3.1 Explainable AI through ILP

Range size, agriculture, altitudinal limits, tail length, tarsus length, and wing length were the most frequently occurring variables across higher confidence rules (≥75%). To enhance clarity and interpretability of our results, we present high-confidence rules that are higher in the rule hierarchy and are less redundant. The complete rule dataset, including non-rounded values, all abnormalities (exceptions to the rule), and the hierarchical order in which rules are applied, is provided in the Supplementary Material.

#### 3.1.1 Purely data-driven models

**With human threats**

- not affected by agriculture → lower risk (94%).
- range size ≤13,500 km² → higher risk (88%).
- tail length ≤304 mm, maximum latitude >1°, range size ≤19 million km², body mass ≤79.2 g → lower risk (78%).
- range size >19 million km² → lower risk (89%).
- tail length >304 mm → higher risk (82%).

**Without human threats**

- range size >15,800 km² → lower risk (88%).
- range size ≤3,270 km² → higher risk (75%).
- beak length ≤18.8 mm → lower risk (79%).

#### 3.1.2 Models informed by data & expert-provided candidate hypotheses

Beyond the five background knowledge rules specified a priori based on scientific literature and Random Forest variable importance (see Methods), the model identified additional high-confidence patterns:

**With human threats**

- not affected by hunting → lower risk (96%).
- occurs in the Indomalayan realm → higher risk (80%).
- maximum latitude >65.5° → lower risk (81%).
- range size ≤5,190 km² → higher risk (75%).
- tail length ≤58.8 mm, generation length >1.95 years → higher risk (75%).
- elevational range >1,275 m, maximum elevation ≤2,200 m → higher risk (76%).

**Without human threats**

- range size >16,400 km² → lower risk (90%).
- beak length ≤18.9 mm → lower risk (76%).
- range size ≤6,440 km² → higher risk (79%).

### 3.2 Random Forest

#### 3.2.1 Random Forest with human threats

Human threats and biogeographical variables were the strongest predictors of extinction risk. Particularly, species affected by agriculture, hunting, invasive species, and with small geographic ranges showed a higher extinction risk. Higher risk was also negatively associated with maximum latitude and elevational range, and positively associated with minimum latitude, generation length, and annual adult survival (Table 2; Figure 1).

**Figure 1.**
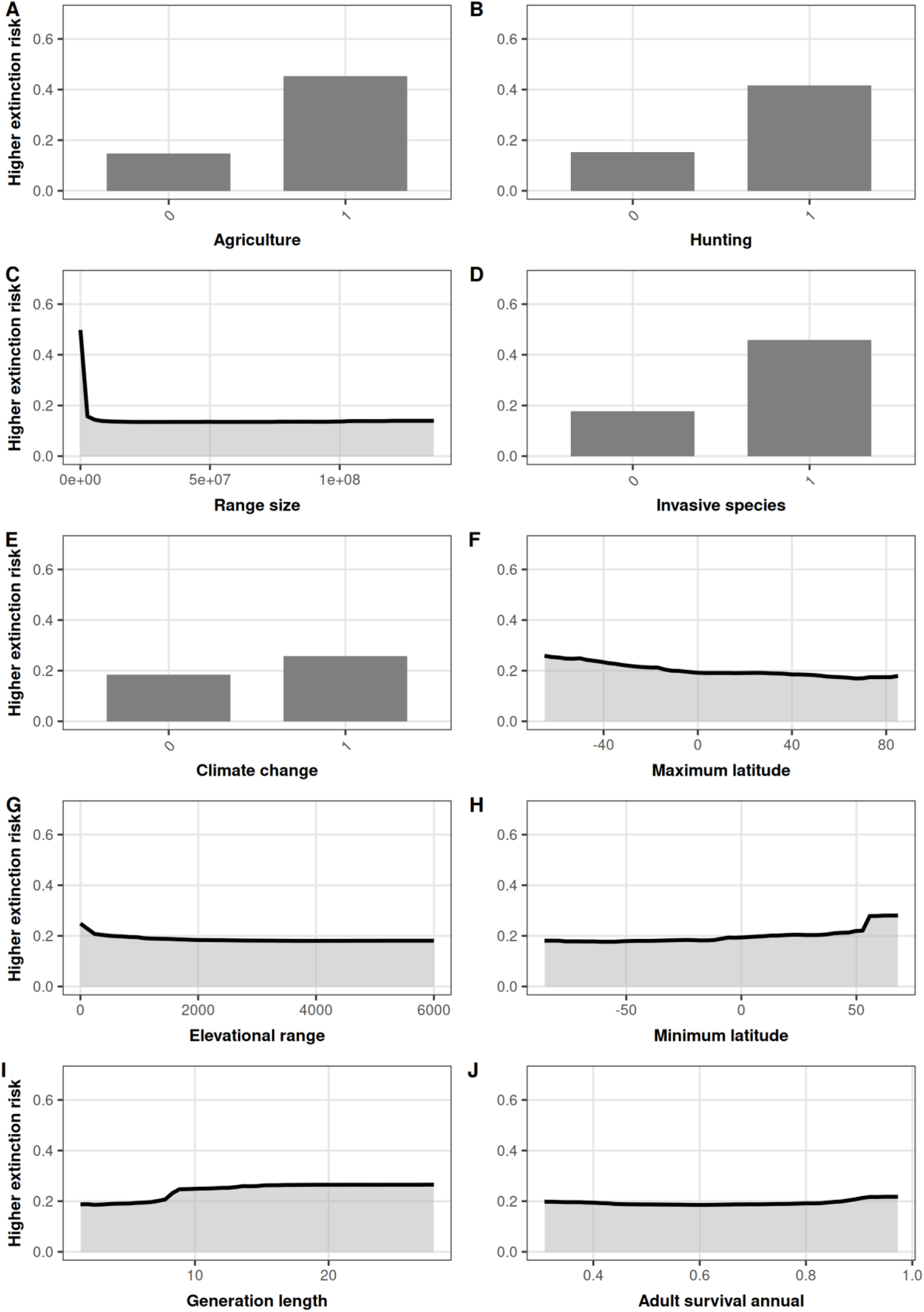
Dependence plots of the ten most important predictors of extinction risk.

#### 3.2.2 Random Forest without human threats

Biogeographic, morphological, and demographic variables were the strongest predictors of extinction risk. Body mass, generation length, maximum latitude and wing length were positively associated with extinction risk, while range size, elevational range, maximum elevation and maximum latitude showed a negative association (Table 3; Figure 2).

**Figure 2.**
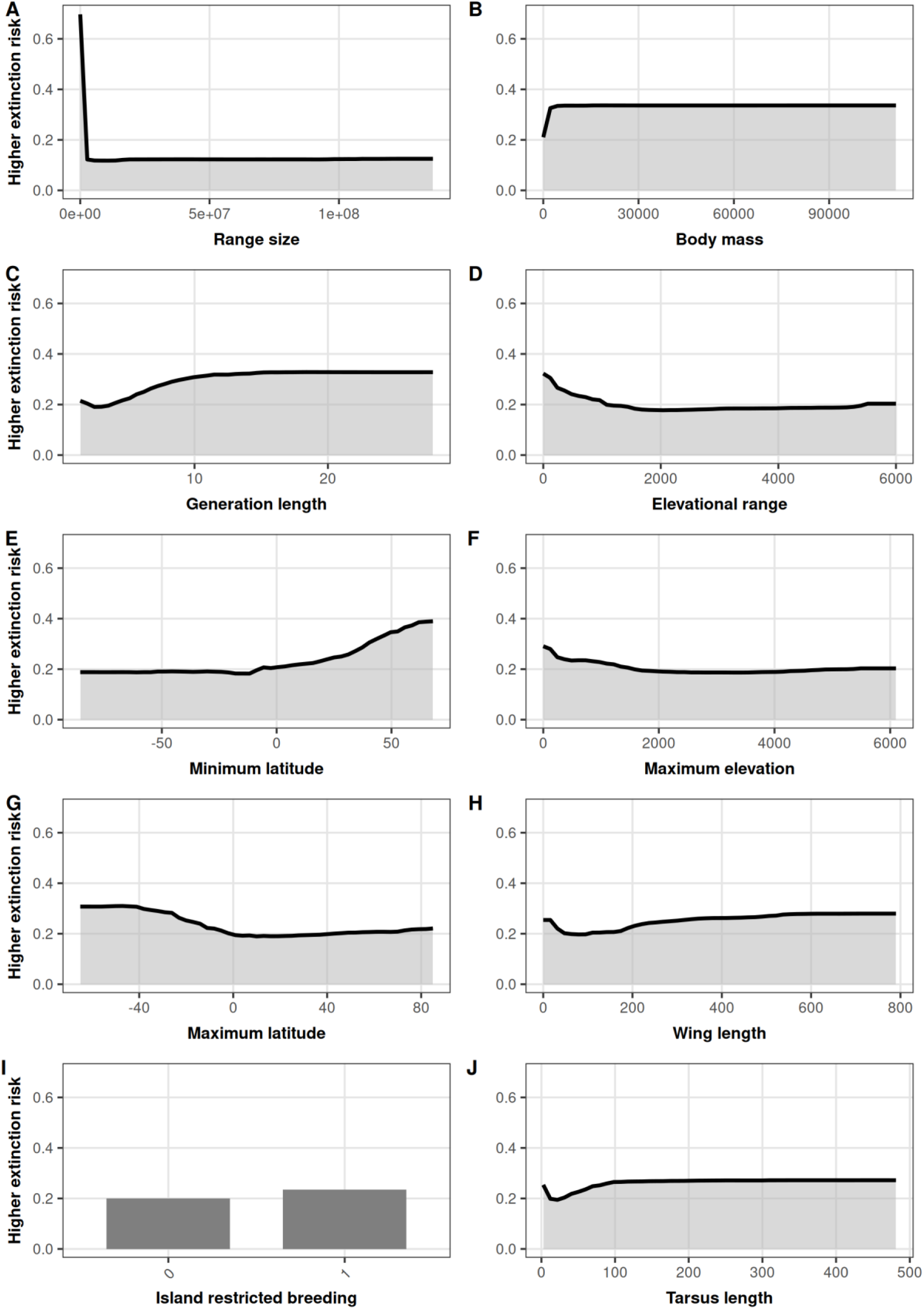
Dependence plots of the ten most important predictors of extinction risk when removing human threats.

## 4. Discussion

Our global analysis encompassing >9,000 bird species reveals a hierarchical and combinatorial structure underlying avian extinction risk. This first fully explainable assessment identified that extinction vulnerability emerges from synergistic interactions among small range size, specific morphological traits, elevational range, and anthropogenic threats, especially agriculture. These findings align with well-established macroecological theory while advancing it by providing explicit quantitative thresholds and rule combinations with associated confidence scores, rather than qualitative associations.

### Quantifiable thresholds of extinction patterns

Explicit numerical thresholds for extinction risk are rarely reported in the literature. One notable exception involves geographic range size. Harris & Pimm (2008), proposed 11,000 km² as a threshold below which species are more likely to be threatened. This is consistent with a large body of evidence showing that small geographic ranges are strongly associated with higher extinction risk, not only for birds but across taxa (Purvis et al. 2000; Böhm et al. 2016), likely reflecting the vulnerability of restricted-range species to stochastic events, or human impacts (González-Suárez et al. 2013). Notably, with 88% confidence, our ILP framework independently identified a very close threshold (13,500 km²). Importantly, when human impact variables were excluded, the extinction threshold shifted to ≤3,270 km² (75% confidence). This indicates that human impacts broaden the set of species classified as high risk to include those with geographic ranges approximately four times larger than those identified as vulnerable based solely on intrinsic traits, constituting strong evidence of how human impacts are shaping extinction paths. These findings also align with the IUCN Red List framework (IUCN Standards and Petitions Committee., 2024), which relies on quantitative range thresholds to assign threat categories. For instance, an Extent of Occurrence below 20,000 km² qualifies a species as Vulnerable under criterion B. Although our range size metric is methodologically distinct from Extent of Occurrence, both approaches indicate that geographic restriction is a leading factor for extinction vulnerability.

Geographical patterns were also evident among rules. We found that species reaching high northern latitudes (maximum latitude >65.5°) were classified as lower risk. This likely reflects the broader geographic ranges characteristic of arctic and subarctic species, as high-latitude species tend to have larger range sizes (Stevens, 1989), which is among the main predictors in our models. Conversely, another rule stated a higher risk for species occurring in the Indomalayan realm (80%). This biogeographical realm, which comprises South, and southeast Asia (Holt et al. 2013), experienced a sharp decline of birds during the 1990s (Butchart et al. 2004) due to intense forest destruction in Indonesia, leading to many species being reassessed and classified under higher risk categories on the IUCN Red List (Butchart et al. 2004).

Indeed, agriculture, heavily related to forest destruction, and the most important variable in our rules, has been categorically linked to avian declines globally (Rigal et al. 2023). Farmland intensification has been particularly associated with declines of insectivorous and ground-nesting birds, largely due to reductions in food resources and nesting habitat (Donald et al., 2001; Benton et al., 2003). Our finding that species not affected by agriculture face substantially lower extinction risk underscores the strong impact of agricultural land-use change on global avifauna. At a global scale, agricultural expansion remains the single largest driver of habitat loss and a leading threat to terrestrial vertebrates, including birds (BirdLife International 2025). This pattern emphasizes the urgent need to integrate biodiversity considerations into agricultural policy frameworks and to promote wildlife-friendly farming practices (Medrano-Vizcaíno et al., 2026).

Hunting also emerged as a significant predictor. While this human pressure may not affect as many species as habitat loss globally, its impacts on targeted species can be catastrophic, particularly for large-bodied species with slow reproductive rates (Ferreiro-Arias et al., 2025; Serrano et al., 2025). Their limited capacity to recover from population declines makes them sensitive not only to hunting, but also to habitat degradation, and other disturbances (Bird et al., 2020).

In this context, previous research has found that large-bodied species are more vulnerable to extinction (Hilbers et al., 2016; Svenning et al., 2024). However, among our rules, body mass alone was not a strong predictor, but its importance was amplified when interacting with other variables. We found that species whose tail length ≤304 mm, maximum latitude >1°, range size ≤19 million km², and body mass ≤79.2 g face lower risk (78%). This rule is consistent with the life history of many small/medium passerines distributed across temperate and subtropical regions and clearly exemplifies the importance of considering synergistic effects of different types of variables when understanding biological processes.

Another rule (complementing logically the previous one) stated that a tail length >304 mm was associated with higher extinction risk (82%). Elongated tails increase body drag and reduce aerodynamic efficiency, impairing flight performance (Zhang et al., 2026). This reduced maneuverability may increase vulnerability to predation and limit the capacity to track resources across disturbed landscapes (Zhang et al., 2026). Moreover, species with long tails tend to be ecological specialists, often associated with structurally complex habitats, whose fitness is disproportionately affected by habitat degradation (Fitzpatrick 1999).

In this line of morphological rules, we also found that a beak length ≤18.8 mm is associated with lower extinction risk (79%). Beak morphology is a key indicator of dietary specialization in birds, with shorter beaks generally associated with generalist diets (Sheard et al., 2020; Tobias et al., 2022). Species with greater dietary flexibility tend to be more resilient to habitat modification and resource availability, being able to shift food sources when preferred items are scarce (Clavel et al., 2011). Conversely, species with elongated beaks are often trophic specialists, whose fitness depends heavily on the availability of specific resources, which can be unavailable due to anthropogenic pressures. Furthermore, smaller species often exhibit faster life histories and higher reproductive rates, and their traits can provide demographic resilience when facing disturbances (Purvis et al. 2000; González-Suárez et al. 2013).

We also found that species occurring at low and mid elevations, despite having broad elevational ranges (>1,275 m), face higher extinction risk (76%) when their maximum elevation does not exceed 2,200 m. This pattern possibly reflects that species unable to reach higher altitudes may be more exposed to human impacts, as upslope habitats tend to be less accessible to agricultural expansion and other anthropogenic pressures (Rahbek et al., 2019). In this sense, a broad elevational range does not necessarily confer resilience if species lack access to more stable, less disturbed refugia at higher elevations. This finding highlights that the protective effect of elevational breadth is contingent on the elevational limits a species can reach.

The quantitative and qualitative convergences of our results with previous research underscore the capacity of ILP to recover biologically meaningful thresholds from complex, high-dimensional data. Our findings have direct implications for conservation prioritization and management. As a large proportion of extinction vulnerability is rooted in species’ biological and biogeographical characteristics, independent of currently documented threats, it would be possible to develop early-warning systems capable of identifying potentially vulnerable species before severe anthropogenic pressures are formally recognized or quantified.

Nevertheless, we identified that the role of human pressures across models was highly relevant. When human threat variables were included, our models achieved substantially higher predictive performance (91% accuracy for CON-FOLD data-driven model vs. 86% without threats). This finding aligns with recent global assessments showing that agriculture affects 73% of threatened bird species, invasive species 42%, hunting 39%, and climate change 37% (BirdLife International 2025).

The synergistic patterns identified by our method emphasize the need for integrated conservation strategies. Recent research also highlighted the importance of an integral approach that combines traits, and human pressures to have a better understanding of extinction risk (Serrano et al., 2025). Similar to their results (based on bayesian hierarchical phylogenetic models), our rules found that particularly, agriculture, hunting, and range size are relevant predictors of extinction. However, it is the interaction among variables, which offers a clearer explanation of extinction patterns. For example, protecting remaining habitat for small-ranged species is necessary but insufficient if human pressures remain unaddressed.

Indeed different morphological traits can maintain distinctive relationships with different human pressures (Stewart et al. 2025). The abatement of hunting and invasive species can reduce the extinction risk for species with long tails and short beaks, whereas the abatement of climate change is associated with a reduced risk for species with short tails and long beaks (Stewart et al. 2025). This suggests that effective conservation interventions need to attenuate multiple human impacts to protect different functional morphologies.

### The advantage of explainable AI for conservation science

Although the overall accuracy of all models was relatively high (>83%), the F1 score, which reflects the balance of precisely predicting lower and higher risk, was lower for Higher risk, particularly in Neural Networks. This results from the unbalanced initial dataset (1,722 Higher risk and 7,331 Lower risk species). The Neural Network can learn a heuristic, something like always classify the result as low risk, since the number of lower risk samples is larger than the number of higher risk species. Conversely, an ILP approach like that used in CON-FOLD alleviates some of the concern that a purely statistical machine learning model may give rise to. Nevertheless, it is clear that the more relevant data we provide to models, the better their performance is. This was evident also in CON-FOLD’s F1 score for Higher risk, which was significantly improved when including human threats, highlighting their importance for assessing extinction.

IUCNN, and similar Neural Network models can predict species classifications from input data but cannot produce verifiable explanations as to what factors, combinations of factors, or values of those factors contribute to a given classification. CON-FOLD, by contrast produces explicit rules with quantitative statements, and confidence scores moving beyond qualitative associations that often rely on subjective interpretations of terms such as “small” or “large”. Our rules contain three key aspects: (1) quantifiable thresholds defining the transition from less to more threatening category, (2) confidence scores that quantify the certainty of the rule, and (3) a combinatorial system that reveals how multiple factors interact to increase or decrease extinction vulnerability. These three aspects together are rarely (likely never) found in conservation science.

The application of explainable artificial intelligence to global bird extinction risk demonstrates that interpretability and predictive performance are not mutually exclusive goals. This AI approach can achieve a high predictive performance equivalent to black box models while maintaining complete interpretability. The performance of the CON-FOLD expert model was better than Neural Networks, and only 2.5 percentage points below Random Forest, representing a minimal trade-off for substantial gains in transparency. This finding challenges the widespread assumption that interpretability necessarily requires sacrificing predictive power (Rudin 2019).

One of the best advantages is the possibility of rule-based outputs to be directly translated into decision criteria. Communicating explicit numerical thresholds to policymakers and practitioners can facilitate actionable conservation planning, bridging the gap between complex analytical models and applied decision-making.

### Limitations and future research

However, there are several considerations for the application of this approach. Using explainable AI in biodiversity research requires not only computational implementation but also ecological literacy in interpreting its outputs. Researchers must recognize that the generated rules are hierarchical, probabilistic statements rather than deterministic laws. Each rule has a specific confidence value and occupies a hierarchical position within the model, which determines its explanatory importance (Mcginness & Baumgartner 2024). A superficial communication of rules without analyzing their confidence levels and hierarchical relationships may lead to overinterpretation. Therefore, proper implementation of this approach demands an integrative understanding of both ecological theory and ILP.

Also, it is important to acknowledge that the accuracy of these models depends on the quality and completeness of the underlying datasets. Geographic range estimates and trait data for certain regions or poorly studied species may be incomplete or biased (Medrano Vizcaíno & Rueda, 2018; Medrano-Vizcaíno & Gutiérrez-Salazar, 2020), potentially affecting the precision of identified thresholds or combinatorial rules. In this context, our focus on birds was partly motivated by the high availability of trait data (e.g., Tobias et al. 2022; Şekercioğlu et al. 2025), which makes this group particularly suitable for large-scale explainable AI analyses. Nevertheless, this methodological framework is transferable across taxa, and the growing availability of comprehensive global trait databases for other groups such as mammals (Soria et al. 2021), amphibians (Oliveira et al. 2017), and reptiles (Oskyrko et al. 2024) creates new opportunities to test whether similar hierarchical and combinatorial extinction patterns emerge across taxa. Our compiled dataset integrating biogeographical, ecological, morphological, demographic, and human-threat variables facilitates future analyses.

As the biodiversity crisis intensifies and multiple human pressures simultaneously threaten an increasing number of species, the need for methodologies that make complex ecological patterns comprehensible becomes paramount. Conservation science does not only require accurate predictions, but also transparent and communicable results that clarify priorities and objectives. In this context, explainable symbolic machine learning frameworks such as recent ILP algorithms constitute a particularly valuable approach for translating complexity into actionable conservation guidance.

## Data availability

The data and code used for ILP (CON-FOLD) and Random Forest are available at: https://github.com/rAIson-Lab/extinction-risk-inference, while the code used for the Neural Network can be found at https://github.com/rAIson-Lab/IUCNN-Modified/blob/master/IUCNN_Runner.ipynb.

