## Supplementary material for "Explainable AI reveals the quantitative hierarchical architecture of global bird extinction risk"

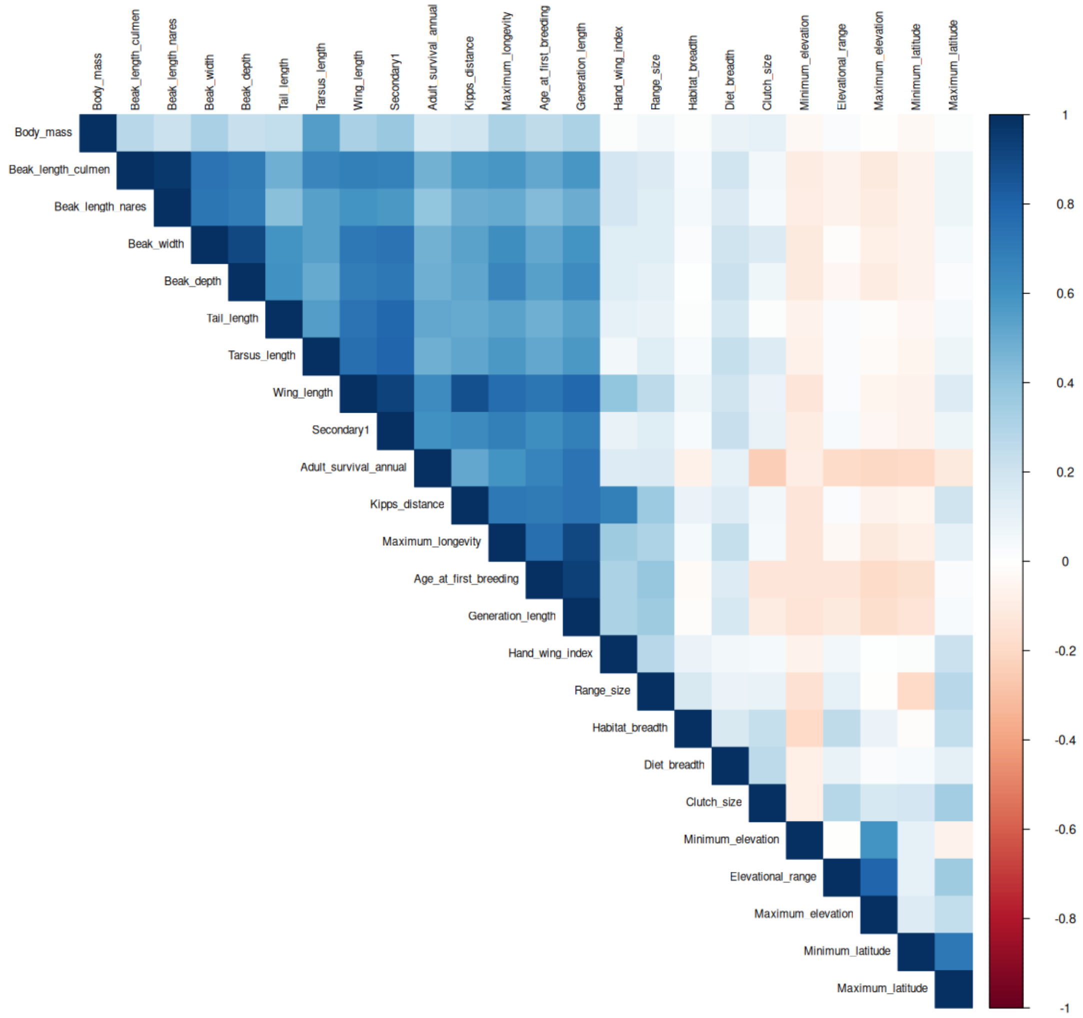


**Figure S1.** Correlation matrix to assess multicollinearity before variable selection

### Complete rules

### Data-driven model with human threats

1. extinction_risk(X,'lower_risk') :- agriculture(X,N2), N2<=0. [confidence: 0.94195]
2. extinction_risk(X,'higher_risk') :- range_size(X,N25), N25<=13551.2. [confidence: 0.87655]
3. extinction_risk(X,'higher_risk') :- not realm(X,'l'), range_size(X,N25), N25<=5136924.07. [confidence: 0.70704]
4. extinction_risk(X,'lower_risk') :- tail_length(X,N11), N11<=304.0, maximum_latitude(X,N13), N13>1.03, range_size(X,N25), N25<=19212039.47, body_mass(X,N26), N26<=79.2. [confidence: 0.77899]
5. extinction_risk(X,'lower_risk') :- range_size(X,N25), N25>19212039.47. [confidence: 0.89024]
6. extinction_risk(X,'higher_risk') :- tail_length(X,N11), N11>304.0. [confidence: 0.82258]
7. extinction_risk(X,'lower_risk') :- not realm(X,'p'), maximum_elevation(X,N22), N22>1600.0, adult_survival_annual(X,N23), N23<=0.87484, range_size(X,N25), N25>354187.36. [confidence: 0.75]
8. extinction_risk(X,'higher_risk') :- not realm(X,'p'), beak_depth(X,N7), N7>3.3, hand_wing_index(X,N10), N10>18.9. [confidence: 0.68155]
9. extinction_risk(X,'lower_risk') :- range_size(X,N25), N25>74565.61. [confidence: 0.66814]
10. extinction_risk(X,'higher_risk') :- minimum_elevation(X,N21), N21<=900. [confidence: 0.69512]
11. extinction_risk(X,'lower_risk') :- range_size(X,N25), N25>17643.08. [confidence: 0.68519]

12. extinction_risk(X,'higher_risk') :- not order(X,'galliformes'). [confidence: 0.625]
13. extinction_risk(X,'higher_risk') :- order(X,'galliformes'). [confidence: 0.55]

### Data-driven model without human threats

1. extinction_risk(X,'lower_risk') :- range_size(X,N21), N21>15801.31. [confidence: 0.8779]
2. extinction_risk(X,'higher_risk') :- range_size(X,N21), N21<=3270.79. [confidence: 0.75379]
3. extinction_risk(X,'lower_risk') :- beak_depth(X,N3), N3<=12.1, minimum_latitude(X,N8), N8>-21.91, island_restricted_breeding(X,N11), N11>0. [confidence: 0.69732]
4. extinction_risk(X,'higher_risk') :- maximum_elevation(X,N18), N18<=2750.0. [confidence: 0.67375]
5. extinction_risk(X,'lower_risk') :- beak_length_culmen(X,N2), N2<=18.8. [confidence: 0.78814]
6. extinction_risk(X,'higher_risk') :- minimum_latitude(X,N8), N8>8.76. [confidence: 0.73529]
7. extinction_risk(X,'lower_risk') :- tail_length(X,N7), N7>95.3. [confidence: 0.65385]
8. extinction_risk(X,'higher_risk') :- wing_length(X,N5), N5>91.3. [confidence: 0.67857]
9. extinction_risk(X,'lower_risk') :- range_size(X,N21), N21>7907.78. [confidence: 0.61765]
10. extinction_risk(X,'higher_risk') :- order(X,'caprimulgiformes'). [confidence: 0.59091]
11. extinction_risk(X,'higher_risk') :- order(X,'passeriformes'). [confidence: 0.55]

### Models informed by data & expert-provided candidate hypotheses with human threats

1. extinction_risk(X,'higher_risk') :- agriculture(X,N2), N2>1, hunting(X,N3), N3>1, invasive_species(X,N4), N4>1, range_size(X,N25), N25<=75321. [confidence: 0.99]
2. extinction_risk(X,'higher_risk') :- agriculture(X,N2), N2>1, hunting(X,N3), N3>1, invasive_species(X,N4), N4>1. [confidence: 0.95]
3. extinction_risk(X,'higher_risk') :- hunting(X,N3), N3>1, invasive_species(X,N4), N4>1, range_size(X,N25), N25<=75321. [confidence: 0.95]
4. extinction_risk(X,'higher_risk') :- agriculture(X,N2), N2>1, hunting(X,N3), N3>1, range_size(X,N25), N25<=75321. [confidence: 0.95]
5. extinction_risk(X,'higher_risk') :- agriculture(X,N2), N2>1, invasive_species(X,N4), N4>1, range_size(X,N25), N25<=75321. [confidence: 0.95]
6. extinction_risk(X,'lower_risk') :- hunting(X,N3), N3<=0. [confidence: 0.95533]
7. extinction_risk(X,'higher_risk') :- realm(X,'i'). [confidence: 0.80256]
8. extinction_risk(X,'lower_risk') :- maximum_latitude(X,N13), N13>65.53. [confidence: 0.80894]
9. extinction_risk(X,'higher_risk') :- climate_change(X,N5), N5>0. [confidence: 0.68644]
10. extinction_risk(X,'lower_risk') :- hand_wing_index(X,N10), N10<=19.0, tail_length(X,N11), N11<=244.5, range_size(X,N25), N25>5194.76, range_size(X,N25), N25<=5280448.27. [confidence: 0.71591]
11. extinction_risk(X,'lower_risk') :- range_size(X,N25), N25>5614395.75. [confidence: 0.69685]
12. extinction_risk(X,'higher_risk') :- range_size(X,N25), N25<=5194.76. [confidence: 0.80682]
13. extinction_risk(X,'higher_risk') :- agriculture(X,N2), N2>0, tarsus_length(X,N8), N8>66.9. [confidence: 0.78889]
14. extinction_risk(X,'lower_risk') :- agriculture(X,N2), N2<=0, generation_length(X,N24), N24<=12.93458. [confidence: 0.6681]
15. extinction_risk(X,'higher_risk') :- tail_length(X,N11), N11<=58.8, generation_length(X,N24), N24>1.9458. [confidence: 0.75]
16. extinction_risk(X,'higher_risk') :- agriculture(X,N2), N2<=0, tail_length(X,N11), N11>61.0. [confidence: 0.7]
17. extinction_risk(X,'lower_risk') :- tail_length(X,N11), N11<=61.0. [confidence: 0.71875]
18. extinction_risk(X,'higher_risk') :- elevational_range(X,N17), N17>1275, maximum_elevation(X,N22), N22<=2200.0. [confidence: 0.75806]
19. extinction_risk(X,'lower_risk') :- maximum_elevation(X,N22), N22>1600.0. [confidence: 0.76087]
20. extinction_risk(X,'higher_risk') :- diet(X,'omnivore'), range_size(X,N25), N25<=2074732.63. [confidence: 0.71875]
21. extinction_risk(X,'lower_risk') :- habitat_breadth(X,N18), N18>2, minimum_elevation(X,N21), N21<=450. [confidence: 0.73214]
22. extinction_risk(X,'higher_risk') :- minimum_latitude(X,N12), N12>-5.71. [confidence: 0.67857]
23. extinction_risk(X,'higher_risk') :- minimum_latitude(X,N12), N12<=-15.03. [confidence: 0.65789]
24. extinction_risk(X,'lower_risk') :- beak_length_culmen(X,N6), N6>20.4. [confidence: 0.67857]
25. extinction_risk(X,'higher_risk') :- not family(X,'sturnidae'). [confidence: 0.59091]
26. extinction_risk(X,'lower_risk') :- order(X,'passeriformes'). [confidence: 0.55]

### Models informed by data & expert-provided candidate hypotheses without human threats

1. extinction_risk(X,'higher_risk') :- elevational_range(X,N13), N13<=800, generation_length(X,N20), N20>4.068, range_size(X,N21), N21<=75321, body_mass(X,N22), N22>130. [confidence: 0.99]
2. extinction_risk(X,'higher_risk') :- elevational_range(X,N13), N13<=800, range_size(X,N21), N21<=75321, body_mass(X,N22), N22>130. [confidence: 0.95]
3. extinction_risk(X,'higher_risk') :- elevational_range(X,N13), N13<=800, generation_length(X,N20), N20>4.068, range_size(X,N21), N21<=75321. [confidence: 0.95]
4. extinction_risk(X,'higher_risk') :- generation_length(X,N20), N20>4.068, range_size(X,N21), N21<=75321, body_mass(X,N22), N22>130. [confidence: 0.95]
5. extinction_risk(X,'higher_risk') :- elevational_range(X,N13), N13<=800, generation_length(X,N20), N20>4.068, body_mass(X,N22), N22>130. [confidence: 0.95]
6. extinction_risk(X,'lower_risk') :- range_size(X,N21), N21>16469.54. [confidence: 0.90208]
7. extinction_risk(X,'higher_risk') :- range_size(X,N21), N21<=3270.79. [confidence: 0.72233]
8. extinction_risk(X,'lower_risk') :- island_restricted_breeding(X,N11), N11>0. [confidence: 0.69326]
9. extinction_risk(X,'higher_risk') :- elevational_range(X,N13), N13<=1000. [confidence: 0.66529]
10. extinction_risk(X,'lower_risk') :- beak_length_culmen(X,N2), N2<=18.9. [confidence: 0.76437]
11. extinction_risk(X,'higher_risk') :- range_size(X,N21), N21<=6446.42. [confidence: 0.78571]
12. extinction_risk(X,'lower_risk') :- maximum_latitude(X,N9), N9>9.76. [confidence: 0.70455]
13. extinction_risk(X,'higher_risk') :- beak_length_culmen(X,N2), N2<=22.7. [confidence: 0.7]
14. extinction_risk(X,'lower_risk') :- beak_depth(X,N3), N3<=8.4. [confidence: 0.625]
15. extinction_risk(X,'higher_risk') :- order(X,'passeriformes'). [confidence: 0.55]
16. extinction_risk(X,'higher_risk') :- order(X,'trogoniformes'). [confidence: 0.55]
